# Deep Learning Allows Assessment of Risk of Metastatic Relapse from Invasive Breast Cancer Histological Slides

**DOI:** 10.1101/2022.11.28.518158

**Authors:** I. Garberis, V. Gaury, C. Saillard, D. Drubay, K. Elgui, B. Schmauch, A. Jaeger, L. Herpin, J. Linhart, M. Sapateiro, F. Bernigole, A. Kamoun, E. Bendjebbar, A. de Lavergne, R. Dubois, M. Auffret, L. Guillou, I. Bousaid, M. Azoulay, J. Lemonnier, M. Sefta, A. Jacquet, A. Sarrazin, J-F Reboud, F. Brulport, J. Dachary, B. Pistilli, S. Delaloge, P. Courtiol, F. André, V. Aubert, M. Lacroix-Triki

**Affiliations:** INSERM U981, Gustave Roussy, Paris-Saclay University, Villejuif, France; Owkin, Paris, France; Oncostat INSERM U1018 labeled Ligue Contre le Cancer, Department of Biostatistics, Gustave Roussy, Paris-Saclay University, Villejuif, France; Department of Pathology, Gustave Roussy, Paris-Saclay University, Villejuif, France; Department of digital transformation and information systems (DTNSI), Gustave Roussy, Villejuif, France; Unicancer R&D, Unicancer, Paris, France; Department of Medical Oncology, Gustave Roussy, Paris-Saclay University, Villejuif, France

**Keywords:** breast cancer, digital pathology, relapse, artificial intelligence, therapeutic de-escalation, outcome prediction

## Abstract

**Background:** Correctly classifying early estrogen receptor-positive and HER2-negative (ER+/HER2) breast cancer (EBC) cases allows to propose an adapted adjuvant systemic treatment strategy. We developed a new AI-based tool to assess the risk of distant relapse at 5 years for ER+/HER2-EBC patients from pathological slides.

**Patients and Methods:** The discovery dataset (GrandTMA) included 1429 ER+/HER2-EBC patients, with long-term follow-up and an available hematoxylin-eosin and saffron (HES) whole slide image (WSI). A Deep Learning (DL) network was trained to predict metastasis free survival (MFS) at five years, based on the HES WSI only (termed RlapsRisk). A combined score was then built using RlapsRisk and well established prognostic factors. A threshold corresponding to a probability of MFS event of 5% at 5 years was applied to dichotomize patients into low or high-risk groups. The external validation, as well as assessment of the additional prognosis value of the DL model beyond standard clinico-pathologic factors were carried out on an independent, prospective cohort (CANTO, NCT01993498) including 889 HES WSI of ER+/HER2-EBC patients.

**Results:** RlapsRisk was an independent prognostic factor of MFS in multivariable analysis adjusted for established clinico-pathological factors (p<0.005 in GrandTMA and CANTO). Combining RlapsRisk score and the clinico-pathological factors improved the prognostic discrimination as compared to the clinico-pathological factors alone (increment of c-index in the validation set 0.80 versus 0.76, +0.04, p-value < 0.005). After dichotomization, the Combined Model showed a higher cumulative sensitivity on the entire population (0.76 vs 0.61) for an equal dynamic specificity (0.76) in comparison with the clinical score alone.

**Conclusions:** Our deep learning model developed on digitized HES slides provided additional prognostic information as compared to current clinico-pathological factors and has the potential of valuably informing the decision making process in the adjuvant setting when combined with current clinico-pathological factors.

## INTRODUCTION

Despite significant progress in classification and treatment over the past two decades, breast cancer (BC) remains the most commonly diagnosed cancer and the leading cause of cancer death for women worldwide (1). Proposing an optimal therapeutic strategy to each patient requires systematic and accurate characterization of each disease. Specifically, estrogen receptor positive (ER+), HER2-negative (HER2-) invasive breast cancer, which accounts for approximately 70% of all invasive BC, and is associated with a wide spectrum of outcomes and treatment requirements. For many of these women a key question remains whether adjuvant chemotherapy with the burden of acute side effects and the potential long-term persistent quality of life (QoL) deterioration (2) can be safely avoided. Furthermore, women with a predicted high risk of metastatic relapse despite current standard treatment are proposed additional strategies based on recent findings, including extended endocrine therapy beyond 5 years, the addition of a CDK4/6 inhibitor, or other strategies (3),(4).

Prognosis definition has been traditionally based on clinical and histopathological factors, such as the patient’s age or the histological grade and subtype of tumors. Biomarker assessment (ER, PR, HER2 and KI67) was added to this estimation and later refined with the inclusion of molecular signatures. Results from the TailorX trial showed that Oncotype Dx, a 21-gene expression molecular test that assesses the 10-year metastasis free survival, could spare up to 85% of women with early BC unnecessary adjuvant chemotherapy, without impacting patient outcomes (5),(6),(7). Currently, several gene expression signatures are endorsed by international guidelines to support clinicians in defining the prognosis of patients with early HR+/HER2-breast cancer and taking adjuvant treatment decisions (8). Furthermore, prognostic tools embedded into publicly available websites have begun to be used as an aid in clinical decision making. Predict Breast, a currently employed online prognostication software, uses known prognostic factors such as tumor size, KI67 index, tumor grade and lymph node status to predict overall survival at 5 and 10 years. However, these methods may be subject to reproducibility and expertise biases (9),(10),(11),(12),(13).

The characterization and prognostication of diseases has evolved over time and recently has expanded to include more sophisticated instruments and methods from the computational field. Artificial intelligence (AI), particularly machine learning (ML) approaches, are increasingly being developed to answer biological and clinical questions. Notably, recent studies have shown the potential of deep learning (DL) models applied to histopathological whole slide images (WSI) to predict patient outcome and unveil features correlated with prognosis in different malignancies, such as brain tumors (14), mesothelioma (15), colorectal cancer (16) and breast cancer (17).

In the era of precision medicine, finding the right treatment for every patient not only requires tools that are accurate, but also that are broadly accessible and able to seamlessly integrate into the clinical workflow.

In this study, we aimed to investigate whether AI applied on tumor WSI could: (i) identify patients who have a substantial risk of metastatic relapse despite receiving standard treatments, and (ii) provide additional prognostic information beyond clinico-pathological prognostic criteria. The ultimate goal was to develop an AI-based digital pathology tool to allow assessment of risk of metastatic relapse.

## MATERIALS, METHODS, DESIGN

### Datasets description

#### GrandTMA

To build our models we used a discovery dataset collected retrospectively from patients treated at Gustave Roussy in France and included in the “GrandTMA” cohort. This cohort comprises all patients newly diagnosed with a breast carcinoma as part of the “One-stop breast clinic” program at Gustave Roussy between October 10th 2005 and February 7th 2013 (18). The inclusion criteria for the present study were: i) diagnosis of invasive breast carcinoma, with or without associated in situ carcinoma, ii) any type of treatment but neoadjuvant chemotherapy, iii) availability of a surgical specimen with a formalin-fixed, paraffin-embedded (FFPE) tumor sample available, iv) complete clinical and therapeutic data, v) follow-up over at least 4 years and updated annually. The exclusion criteria were: i) exclusive non-invasive tumors, ii) cytology-only available cases, iii) absence of follow-up, iv) other non-adenocarcinomatous lesions of the breast. This led to the inclusion of 1802 patients diagnosed with early invasive BC (1429 ER+/HER2-, 110 ER+/HER2+, 70 ER-/HER2+, 193 ER-/HER2-), with at least 1 available hematoxylin-eosin-saffron (HES)-stained tumor slide from the surgical specimen at the Pathology Department (details in Supplementary Figure 1). Biomarker status (ER, PR, and Ki67 immunohistochemistry (IHC) expression, and HER2 protein expression/gene amplification) was defined and determined locally according to the current recommendations of the College of American Pathologists and the American Society of Clinical Oncology (CAP/ASCO (19),(20)), and the French recommendations of the study group on immunohistochemical prognostic factors in breast cancer (GEFPICS (21)). ER and PR expression positivity was defined as an IHC staining of at least 10% of tumor cells, as is standard in several European countries. Lesions were considered positive for HER2 (score 3+) if the number of tumor cells with a complete and intense membrane IHC staining exceeded 10% of the whole infiltrating tumor cells population; equivocal (score 2+) if the number of tumor cells showing a complete and moderately intense membrane IHC staining, or an incomplete basolateral membrane IHC staining of moderate to severe intensity exceeded 10% of the total infiltrating tumor population; and negative in the remaining cases (scores 0 and 1+). When dichotomized, Ki67 cut-off was defined according to the current recommendations (KI67 = 20) (20), (22).

Image acquisition was performed using an Olympus VS120 slide scanner at 20x magnification. In order to avoid scanning issues that might affect subsequent image analysis, all slides were checked by a pathologist after digitization to discard slides with insufficient quality and rescanned when necessary (blurred images, need of slides re-mounting when coverslip was damaged).

This work was carried out in accordance with the provisions of the Public Health Code applicable to research not involving the human person (Public Health Code - Article R1121-1 amended by Decree no. 2017-884, May 9^th^, 2017) and therefore it does not come under the jurisdiction of a Committee for the Protection of Persons. It obtained the favorable opinion of the expert Committee for Research, Studies and Evaluations in the field of breast pathology, as well as of the Ethics Committee (Data Protection Office, Gustave Roussy). It has been submitted to the National Commission for Computing and Liberties (CNIL) under reference N° F20220121170839 and has been declared in accordance with the reference methodology MR-004. The patients involved were informed of the research via an information letter distributed by post mail with the possibility of opposing the study.

#### CANTO

For the purpose of an external validation of our model, we used a dataset from the French observational and prospective CANTO cohort (NCT01993498) (23). In this cohort, patients were included at diagnosis of their invasive breast cancer, before any treatment, following the given criteria: i) women only, ii) aged over 45 years old, iii) HER2- and ER+ (same definition as for the GrandTMA cohort), iv) with a histologically invasive breast cancer diagnosed, v) with no clinical evidence of metastasis at the time of inclusion. Out of 14,000 patients accrued in CANTO so far, 1705 ER+/HER2-EBC patients had a minimum follow-up of 5 years and were eligible for the present study (708 patients from Gustave Roussy and 997 patients from other cancer centers of the UNICANCER group). None of these patients were also included in the GrandTMA cohort. Thirty-three patients from Gustave Roussy had incomplete data and were excluded from the study, resulting in 675 HES slides available with clinical data. From the other centers of the CANTO cohort, we had access to 214 HES slides from resection used for primary diagnosis (details in Supplementary Figure 2). In total 889 patients had exploitable HES slides, together with full clinico-pathological features (described in Supplementary Table 1).

All data collections were performed in the framework of the CANTO clinical trial (NCT01993498), in compliance with all legal requirements. All patients included in the study were informed through the website https://mesdonnees.unicancer.fr/ on the reuse of their data for a separate objective with the possibility of opposing the study.

#### Endpoint

The chosen endpoint for survival data analysis was metastasis-free survival (MFS) at five years, defined as the time from initial surgery to occurrence of metastatic event or death before five years. Local relapse or axillary lymph node recurrence events were ignored. Patient’s follow-up was censored at the time of contralateral breast cancer, second non-breast primary cancer or last available date of follow-up.

### Model Description

To develop our risk score from histology slides we used a method composed of three steps: i) tissue tiling, ii) feature extraction, iii) creation of a risk score. The transformation of the score into a probability of occurrence of a MFS event before five years and the selection of a threshold are two additional steps, detailed in Statistical Analysis.

#### Tissue segmentation and tiling

Each of the Whole Slide Images (WSI) was first divided into small squares, 76 × 76 micrometers in size (224 × 224 pixels) called “tiles”. This tiling was performed by first segmenting the tissue, using a pre-trained U-Net neural network (24) that discarded the background, and artifacts of scanning or preparation. This segmented tissue was then divided into N (ranging from 10,000 to 75,000) tiles.

#### Feature Extraction

The N tiles were embedded into D-dimensional feature vectors using a pre-trained CNN (Fig. 1A). We implemented Momentum Contrast v2 (25), a self-supervised learning algorithm that improved performance for various prediction tasks in previous studies (26) trained on the Cancer Genome Atlas Colon Adenocarcinoma (TCGA-COAD) dataset (27)). Multiple data augmentations were applied while the model was optimized for 200 epochs (approximately 30 hours) on 16 NVIDIA Tesla V100 Graphics processing units (GPU). This frozen pre-trained algorithm was then used to extract features during training and inference.

**Figure 1:**
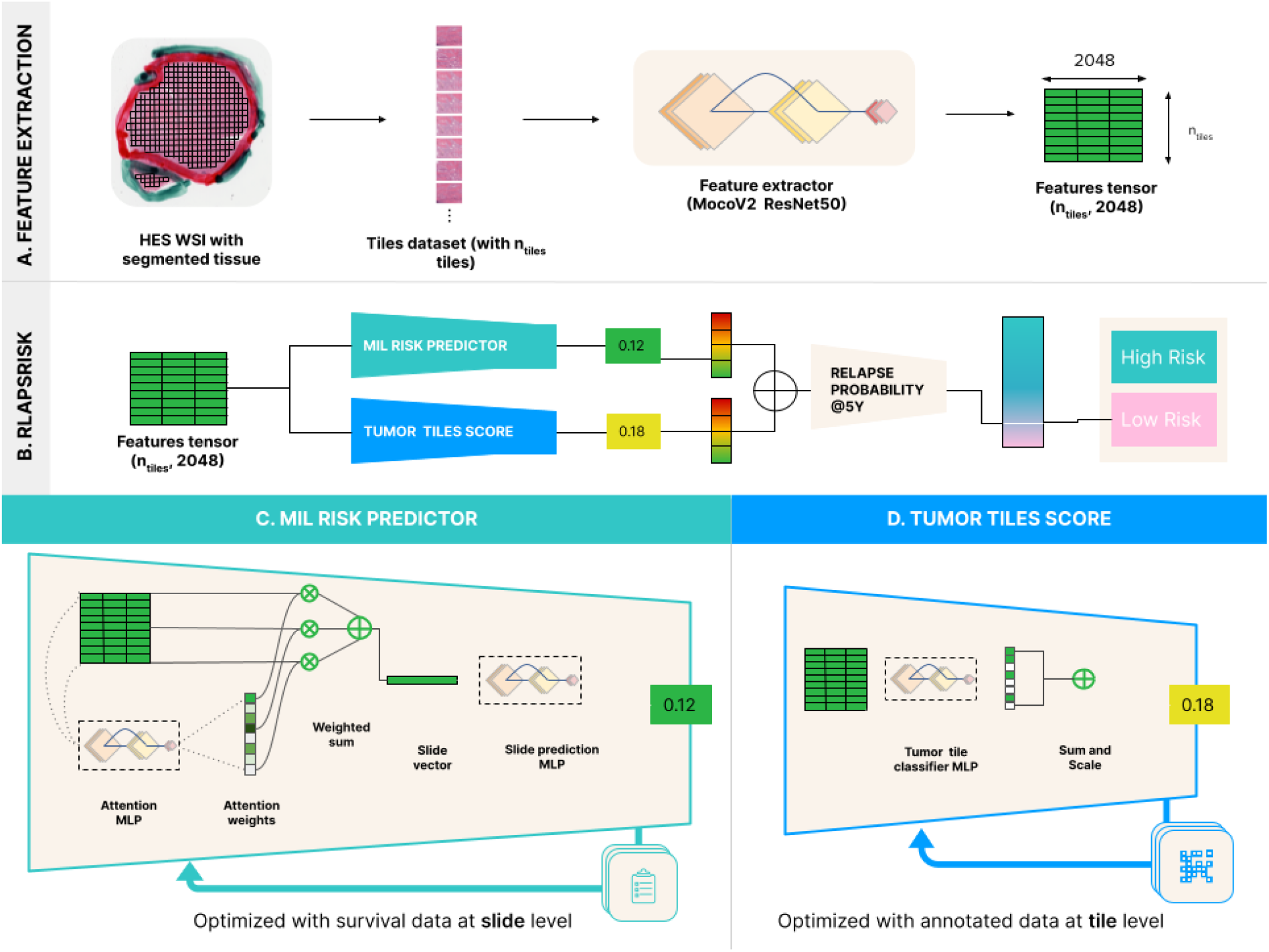
RlapsRisk algorithm overview

#### Risk prediction using Multiple instance learning

The N feature vectors were then aggregated using a multiple instance learning (MIL) model trained to predict MFS at five years (Fig. 1C). We reimplemented the attention based model called DeepMIL proposed by Ilse et al. (28). A linear layer with L neurons (L = 256 here) was applied to the embedded features followed by a Gated Attention layer with L hidden neurons. A multilayer perceptron (MLP) with 128 neurons was then applied to the output. To speed-up training, only a random subset of 8000 tiles per WSI was used, while all tiles of a slide are processed for inference. DeepMIL was trained using an extension of the standard cross-entropy loss used to train survival prediction models with right-censored data (29).

#### Integrating a tumor-related feature in the algorithm

Our preliminary analyses highlighted that the number of tumor tiles contained in each slide was associated with distant relapse and yet was not captured by the model. We therefore incorporated this feature by classifying all the tiles of a slide as tumor vs non-tumor (Fig. 1D). An ensemble of four MLPs trained in a patch-based supervised learning approach was used.

The combination of the predicted DeepMIL risk score and the tumor count score was done by scaling and summing both features, forming the RlapsRisk score (Fig. 1B).

#### Method for interpretability features assessment

Training an AI model on digital slides from diagnosis to predict metastatic relapse is an original approach compared to recent research works in Digital Pathology, that generally predict well known pathological features (e.g. histological grading or KI67 index). However, bypassing human knowledge in the training phase requires even more explanations of the functioning of the model. We detail herein our method to identify and characterize typical areas on a well defined set of slides that had extreme risk scores. Interpretability relies on the possibility to access the relevant information for our model that is contained in input data or learned by the model itself.

In the model, each tile was associated with an attention score that was used in the final weighted average to obtain the input vector for the risk predictor (see Fig. 1). However, this score did not provide information about the impact on prognosis of the tile. To overcome this limitation, we computed the Shapley value (30) associated with each tile, which measures the marginal contribution of each tile to the overall risk score to assess its positive or negative contribution to the final risk score predicted by the final MLP.

For 20 slides of the validation dataset (10 classified with highest RlapsRisk scores, 10 classified with lowest RlapsRisk scores by our model), we computed the Shapley values of each tile and extracted those with the 10 highest computed contributions (in the case of the highest RlapsRisk scores, those tiles constituting the high risk contribution group) and 10 lowest ones (in the case of the lowest RlapsRisk scores, constituting the low risk contribution group) for each slide. Those tiles were further reviewed and annotated by two expert pathologists blinded to the predicted outcome.

Fifty-six histological features were recorded, encompassing tumor architecture patterns, stroma features, presence of different cell types and tiles’ localization. Only twenty-three histological features were kept in the analysis after excluding those with low agreement between the two experts (Cohen’s kappa < 0.21, (31)). The proportions of appearance of each feature in the highest and lowest contribution groups were compared with a two proportion Z-test, statistical significance was calculated using the Bonferroni adjustment, resulting in a corrected alpha value of 1E-3.

### Statistical Analyses

The primary objective was to evaluate the additional 5-years MFS prognostic value of RlapsRisk Score to the current clinico-pathological criteria in patients with ER+/HER2-early breast cancer. The secondary objectives consisted in evaluating the capacity of a model combining clinico-pathological criteria and RlapsRisk to better dichotomize patients between high risk and low risk of developing 5-years MFS events than a model based on clinical factors only. This comparison was assessed on all population and in different subgroups of clinical interest (histological grade 2, clinical intermediate risk of relapse defined in Table 1, patient pre-versus post-menopause, with or without lymph node invasion, treated with or without adjuvant chemotherapy).

**Table 1:** Clinical risk groups definition

| High Risk<br>(At least one criteria) | Intermediate | Low Risk<br>(All Criteria) |
| --- | --- | --- |
| • pT3<br>• pN1 & pre-menopause<br>• pN2 & post-menopause<br>• Grade 3 | Other Situation | • pT1<br>• pN0/i+/mi<br>• Grade 1<br>• KI67<15-20%<br>• No lymphovascular emboli |

We first developed a clinical score to predict the 5-year MFS from a multivariable Cox model trained using the discovery dataset (hereafter the Clinical Score). We considered this score as our baseline reference that we compared to a score based on a Cox Model adjusted for the clinical factors and RlapsRisk score to assess the additional predictive value of the RlapsRisk score to the standard clinical factors (hereafter the Combined Score). Adjuvant treatment regimen was not a prognostic factor in multivariate analysis (both with or without RlapsRisk score) and was therefore removed from all prognostic models.

The discrimination power of these two scores was first compared using C-index (32) on the discovery dataset with five stratified folds cross-validation strategy, with three repeats. Due to the small number of events, we stratified this cross-validation on the events to preserve a minimal number of events per fold. Data from the CANTO cohort were held out from the training series and were used only for a one-shot external validation and the assessment of the discrimination of each model.

For each risk score, we fitted a Weibull AFT (Accelerated Failure Time) model on the discovery dataset to transform risk scores into probabilities of occurrence of MFS event before 5 years. This step was used to identify the threshold of each risk score corresponding to a probability of 5-years MFS event of 5% defined by the Weibull Models. This rate corresponds to the 5-year MFS interpolation using an exponential model from the 10-year MFS of 10%, which is the output of the molecular signatures currently used in clinical practice (33).

Thereby defined, the thresholds were applied to each risk score in order to dichotomize patients between high-risk and low-risk categories. When dichotomized, the scores will be referred to as Classifiers (e.g. Combined Model Classifier).

We performed Kaplan-Meier analyses to assess the association of each classifier with the MFS (the presented HR and their 95%CI corresponding to the related univariable Cox models). In order to evaluate the prediction performance of these classifiers in term of discrimination of the risks of events before five years, we also computed the cumulative sensitivity as well as the dynamic specificity that are natural extensions of the so-called sensitivity/specificity to the particular setting of time-to-event outcomes that may be censored (34).

Following that same methodology, we also investigated those performances within subgroups of patients known to be of intermediate prognosis such as the histological grade 2 population and the “clinical risk group” defined by the table 1.

We also assessed the performance of our tool across groups of patients at different risk of recurrence, as defined according to known prognostic factors, such as pre-menopausal patients vs. post-, patients with lymph node invasion vs. without, patients treated with chemo-endocrine therapy vs. endocrine therapy alone.

Performance assessment though Harrell’s c-index and Kaplan-Meier analyses were performed with uni- and multivariate Cox proportional hazards models implemented in the lifelines (0.27.4) package of Python, cumulative sensitivities and dynamic specificities were computed using the scikit-survival package (0.19.0).

## RESULTS

### RlapsRisk Score and Prognosis

Additional information on each cohort is detailed in supplementary Table 1. On both datasets, RlapsRisk score was an independent prognostic factor of MFS (Table 2) when integrated into a multivariable Cox model including histological grade, age, lymph node invasion, tumor size, and Ki67 expression. A Cox model using the Histological Grade as a continuous variable was compared using C-index to a Cox Model using a dummy encoded version of that same variable. Both models had equivalent performances, so we decided to keep the Grade as a continuous variable. All the other variables were considered continuous.

**Table 2A:** Multivariable Cox proportional hazard models for estimating the contribution of prognostic variables on MFS on GrandTMA

| Grand TMA | Variable | Cox Model without RlapsRisk |  | Cox Model with RlapsRisk |  |
| --- | --- | --- | --- | --- | --- |
|  |  | Unit HR (95% CI) | P | Unit HR (95% CI) | P |
| Multivariable<br>Cox model | Histological Grade | 1.26 (0.92–1.73) | 0.15 | 1.38 (0.99–1.92) | 0.22 |
|  | Age | 1.37 (1.12–1.67) | <0.005 | 1.26 (0.93–1.54) | 0.02 |
|  | Lymph Node Invasion | 1.12 (1.08–1.15) | <0.005 | 1.12 (1.08–1.15) | <0.005 |
|  | Tumor Size | 1.32 (1.16–1.51) | <0.005 | 1.25 (1.08–1.44) | <0.005 |
|  | Ki67 expression | 1.44 (1.21–1.71) | <0.005 | 1.39 (1.16–1.66) | <0.005 |
|  | RlapsRisk score | N.A. | N.A. | 1.27 (1.13–1.43) | <0.005 |
|  | C-index (cross-validation) | 0.78 |  | 0.80 |  |

**Table 2B:** Multivariable Cox proportional hazard models for estimating the contribution of prognostic variables on MFS on CANTO *C-index were computed from the Cox multivariable models trained on Grand TMA.

| CANTO | Variable | Cox Model without RlapsRisk |  | Cox Model with RlapsRisk |  |
| --- | --- | --- | --- | --- | --- |
|  |  | Unit HR (95% CI) | P | Unit HR (95% CI) | P |
| Multivariable<br>Cox model | Histological Grade | 1.84 (1.16–2.91) | 0.01 | 1.61 (0.94–2.76) | 0.08 |
|  | Age | 1.11(0.85–1.44) | 0.46 | 1.15 (0.87–1.53) | 0.33 |
|  | Lymph Node Invasion | 1.15 (1.07–1.23) | <0.005 | 1.11 (1.03–1.20) | 0.01 |
|  | Tumor Size | 1.26 (1.05–1.50) | 0.01 | 1.16 (0.93–1.46) | 0.19 |
|  | Ki67 expression | 1.21(1.00–1.47) | 0.05 | 1.23 (1.01–1.51) | 0.04 |
|  | RlapsRisk score | N.A | N.A | 1.38 (1.12–1.71) | <0.005 |
|  | C-index (validation)* | 0.76 |  | 0.80 |  |

On the validation dataset, our results highlighted an increase of the model discrimination adding RlapsRisk to the clinical factors, with a C-index of 0.80 compared to the clinical factors alone (C-index = 0.76, i.e. +0.04, p-value <0.005).

We then assessed the prognostic performance of RlapsRisk score and the clinical factors in patient subgroups defined by standard clinico-pathological factors, by the clinical risk groups (Table 1) and by adjuvant treatment regimen. The assessment of potential heterogeneities in these subgroups was conducted by Cox regression analyses. When a factor was used to build a subgroup it was removed from the associated Cox multivariable model.

A higher RlapsRisk score was associated with an increased risk of distant recurrence in all subgroups except for patients with pN=2, pN=3 and pT=3 (see Figure 2).

**Figure 2:**
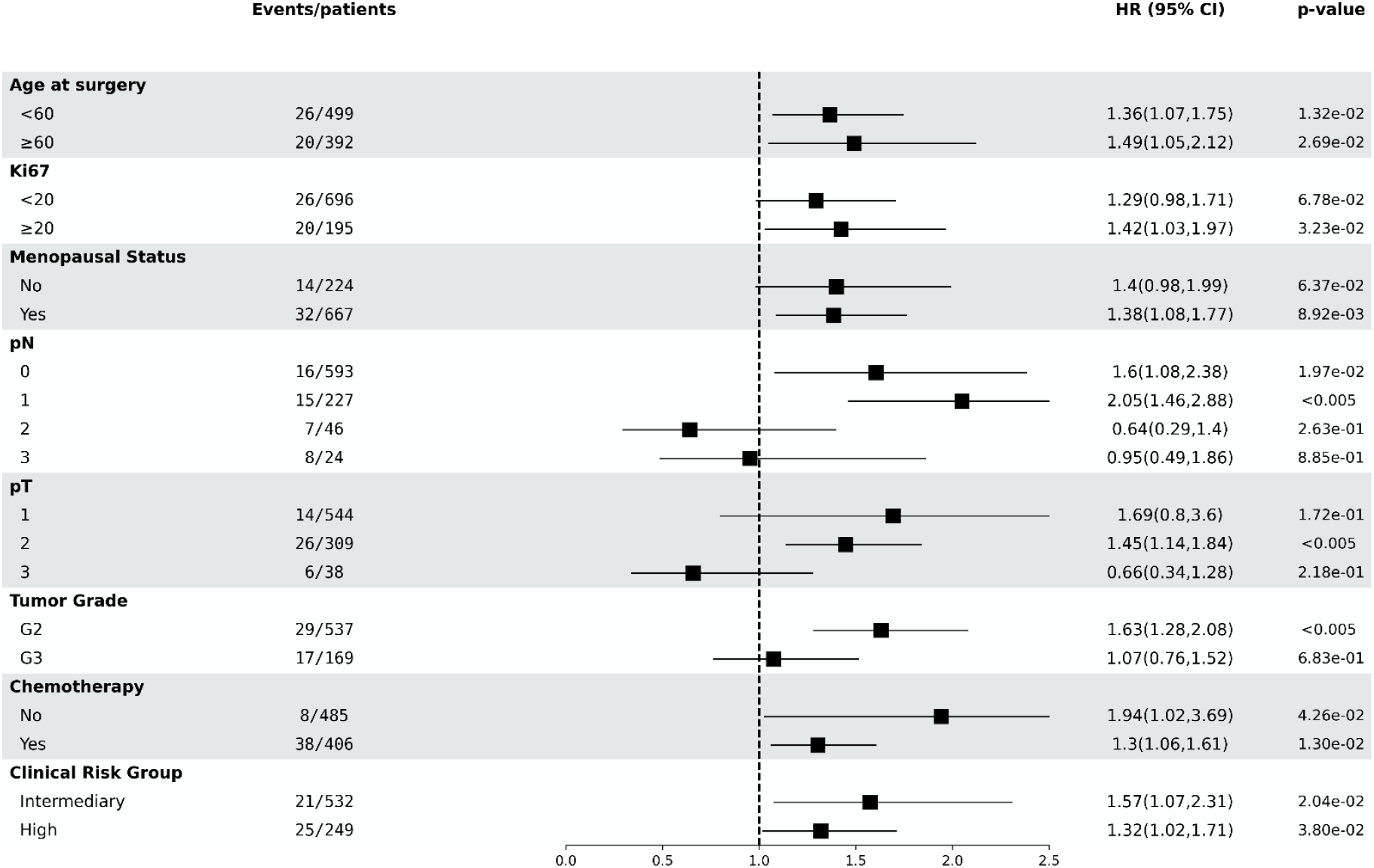
Forest plot of the adjusted RlapsRisk Score HRs. Each square of the forest plot represents the HR of the adjusted RlapsRisk Score (as continuous variable) and clinical factors in the subgroup of patients defined by the variable category in the first column of the table. The HRs 95% CIs were represented by the horizontal lines. Tumor Grade 1 and Low Clinical Risk Group were removed as no MFS events were recorded in those subgroups.

### Clinical validity of RlapsRisk

After applying the stratification of population according to the classifiers, Kaplan–Meier analyses showed significant differences in distant recurrence between low-risk and high-risk patients on the discovery and on the validation dataset, as summarized on Table 3 and highlighted on Figure 3.

**Table 3:** Classification of Patients according to each classifier * probability of 5-years MFS event inferior to 5% ** probability of 5-years MFS event superior to 5%

|  | CANTO |  |  |
| --- | --- | --- | --- |
|  | RlapsRisk Classifier | Clinical Score Classifier | Combined Model Classifier |
| **Low Risk\*** | 562 | 663 | 658 |
| **High Risk\*\*** | 327 | 226 | 231 |
| **% of patients with MFS events in the Low Risk group** | 1,42% | 1,96% | 1,22% |
| **% of patients with MFS events in the High Risk group** | 7,95% | 9,29% | 11,26% |
| **HR** | 4.36 | 4.25 | 6.99 |
| **CI** | 2.32–8.18 | 2.36-7.64 | 3.73-13.09 |
| **p-value** | <0.005 | <0.005 | <0.005 |

**Figure 3:**
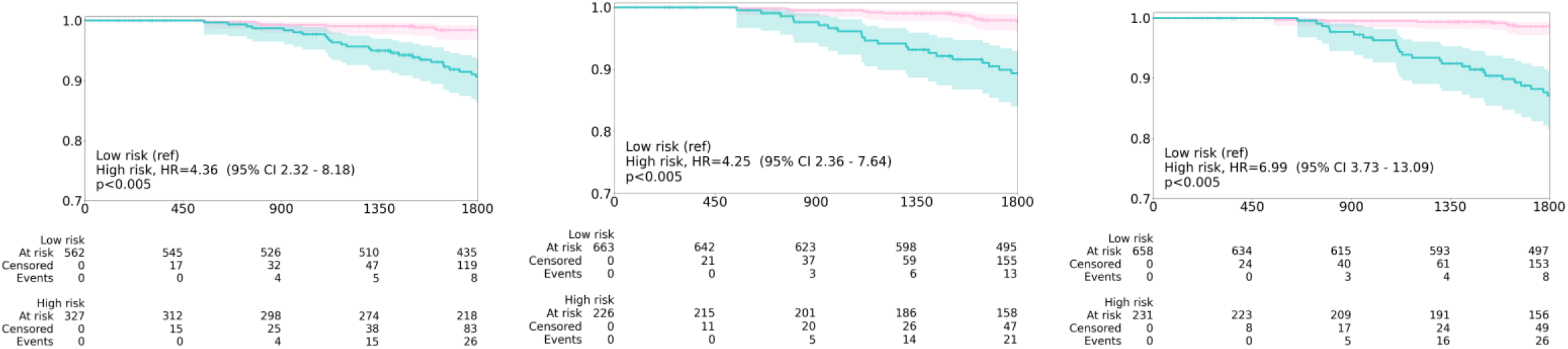
Stratification performed by RlapsRisk Classifier (left) Clinical Score Classifier (middle) Combined Model Classifier (right) of patients from the Canto cohort.

The discriminative power of the Model Combined Classifier on all presented subgroups (Table 4) suggest that when combined with the current clinical factors RlapsRisk could be used as an additional layer of information for better defining the risk of recurrence of each patient.

**Table 4:** Performance of the Classifiers in different subgroups of the CANTO cohort (external validation)

| Subgroup | N | RlapsRisk Classifier |  | Clinical Score Classifier |  | Model Combined Classifier |  |
| --- | --- | --- | --- | --- | --- | --- | --- |
|  |  | Cumulative Sensitivity | Dynamic Specificity | Cumulative Sensitivity | Dynamic Specificity | Cumulative Sensitivity | Dynamic Specificity |
| **All Population** | 889 | 0.77 | 0.67 | 0.61 | 0.76 | 0.76 | 0.76 |
| **Grade 2** | 534 | 0.80 | 0.68 | 0.46 | 0.79 | 0.74 | 0.79 |
| **Intermediate clinical risk group** | 529 | 0.84 | 0.73 | 0.41 | 0.82 | 0.67 | 0.84 |
| **Lymph Node Invasion** | 296 | 0.71 | 0.5 | 0.47 | 0.70 | 0.8 | 0.56 |
| **No Lymph Node Invasion** | 593 | 0.90 | 0.74 | 0.38 | 0.82 | 0.69 | 0.85 |
| **Pre-Menopause** | 223 | 0.81 | 0.57 | 0.50 | 0.83 | 0.72 | 0.79 |
| **Post-Menopause** | 666 | 0.75 | 0.70 | 0.66 | 0.73 | 0.78 | 0.75 |
| **adjuvant CT** | 405 | 0.75 | 0.48 | 0.66 | 0.60 | 0.78 | 0.57 |
| **no adjuvant CT** | 484 | 0.85 | 0.80 | 0.41 | 0.88 | 0.70 | 0.91 |

On the entire dataset, with a same dynamic specificity of 0.76, the Model Combined Classifier has a gain of 15 points in cumulative sensitivity in comparison to the Clinical Score Classifier alone. When looking at the Histological Grade 2 group and the “intermediate clinical risk” group (defined in Table 1), with a Dynamic Specificity equivalent and a gain of 18 and 26 respectively in Cumulative Sensitivity, the Model Combined Classifier largely outperforms the Clinical Score Classifier. These figures expose a benefit of adding RlapsRisk score to the current clinical factors for a better estimation of prognosis within subgroups with a difficult prognosis estimation.

Notably, the combination of the two models is the only model achieving a stable performance for both pre and post menopausal patients.

Additional exploratory Kaplan–Meier analyses highlighted significant differences between high-risk and low-risk in all these subgroups (see supplementary material Figures 1 to 5).

### Model interpretability: features assessment

The most predictive features of high risk of relapse according to RlapsRisk Classifier (Fig. 4B) were the presence of high tumor cell content (P < 0.0001), high degree of nuclear pleomorphism (P < 0.0001), massive architecture (P < 0.0001) with low tubule formation (P < 0.0001) and trabecular structures (P < 0.0001) as well as mitotic activity (P < 0.0001). Multiple features were associated with low risk of relapse such as fibrosis (P < 0.0001), presence of vascular structures (P < 0.0001) and isolated tumoral cells (P = 1.2E-03). Interestingly, tiles associated with low risk (Fig. 4A) were not only located in tumor stroma (P < 0.0001), but also in the adjacent normal tissue (P < 0.0001). Moreover, the model was also able to capture complex cell interactions, as spatial mixing of fibroblasts and tumor infiltrating lymphocytes (TILs) were also retrieved in low risk tiles (P = 2.5E-04), a feature of good prognosis highlighted in a recent study (35). These results showed that our model was able to capture well-established prognostic factors as well as complex and novel patterns to predict patient outcomes.

**Figure 4:**
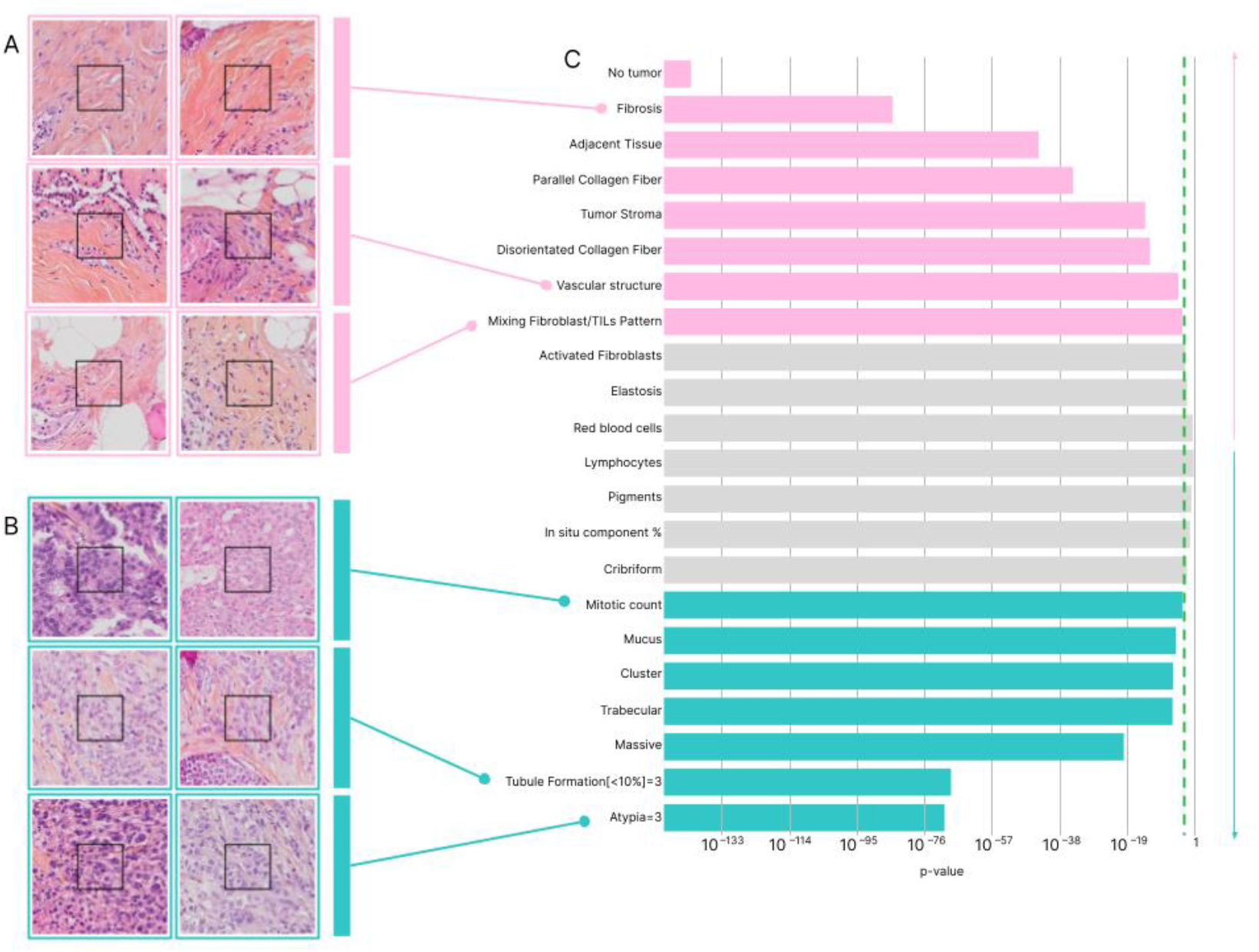
Interpretability

## DISCUSSION

Digital pathology integration is increasing in everyday practice, and workflows that include digitization of glass slides are no longer an exception in pathology laboratories. This ongoing transition is paving the way for the implementation of AI-based digital pathology medical devices in clinical practice.

In this study, we designed a new digital pathology test, which predicts distant recurrence at 5 years after surgery in early-stage breast cancer, with ER-positive and HER2-negative status. To provide a risk score, the method uses just a simple, standard-stained and scanned tumor slide already available for diagnostic purposes in the pathology laboratory.

RlapsRisk score had prognostic value independent to established clinicopathologic factors and provided additional information to the clinical factors. When combined together, clinical factors and RlapsRisk were able to dichotomize patients into low-risk and high-risk groups with a strong discriminative power, on the entire population, but also in subgroups of interest.

Contrary to other digital pathology approaches predicting morphological features, such as histological grade or KI67 index that are ultimately indirectly linked to prognosis (36),(37),(38),(39), we trained our AI model to directly predict the MFS. Not only did we not require local annotations to train the algorithm, which takes as input the entire WSI, but our prediction task was directly addressing our clinical question, i.e. prediction of 5-years MFS.

A different approach presented by Wang et al (40) attained a risk prediction from Nottingham histological grade through the re-stratification of the intermediate category (NHG2). NHG2, which encompassed the largest group of patients, was dichotomized in a low-grade subgroup and a high-grade subgroup, achieving a HR of 2.94 (95% CI 1.24-6.97, P = 0.015) for the stratification between the two groups according to Recurrence Free Survival (RFS). While the presented tool had an important prognostic implication, it was centered on the intermediate risk group and it addressed the question of recurrence risk in an indirect manner (40). However, the strength of their approach is served by the straightforward explicability of the model. Highlights of simple interpretability features will ensure the pathologist that a given classification was performed according to expected criteria: tubule formation, nuclear pleomorphism and mitotic activity, in the case of histological grading. Similarly, in our study the validation cohort RlapsRisk Classifier achieved a high discriminative power on NHG2 patients with a sensitivity of 0.8 and a specificity 0.68, respectively, and illustrated by an HR of 4.15 (95% CI 1.88-9.15, P < 0.005) for MFS (see supplementary material) although the use of a different endpoint which excludes locoregional relapse, contralateral tumors and death contrary to RFS.

Interpretability is indeed a compelling concern for the use of AI in digital pathology, even presented as a crucial challenge by Duggento et al (41). In our study, we aimed to investigate the features that supported a low-risk or high-risk prediction from RlapsRisk. Analyzing the most predicting tiles, we confirmed that the model retrieves well-established criteria alongside complex and novel histological patterns that have an impact on patient outcomes, such as high atypia or mitotic activity, without having to use any pre-annotated data. In this manner, we propose a natively interpretable model to step out of the black box paradigm, a major hurdle toward a broad adoption of any AI solution in clinical practice to help clinicians tackle medical challenges (42).

Currently, one of these challenges in ER+/HER2-EBC management is the adaptation of the treatment according to the risk of the patient. Reducing the number of unnecessary adjuvant chemotherapies or even endocrine therapies to improve quality of life while maintaining an equivalent survival rate of the patients is the key challenge. This group is currently the target population for molecular signatures, where the issue is to use the genomic score to decide on an adjuvant chemotherapy or to decide on a de-escalation in certain patients. However, the precise conditions of these molecular tests restrict their indication, their use is not generalized and they are not covered by the health care system in many countries (43). These facts expose a gap where a simpler, less-expensive, routinely available tool, could support clarity in decision-making, replace molecular signatures or, at least, work as a pre-screening test for ulterior indication of onerous molecular determinations on selected cases. This approach has already been explored in some studies (44), and a comparison study between molecular signatures and RlapsRisk in a similar setting to the TRANSATAC study (45) would provide helpful insights to that matter.

Even though our study presents an innovative and useful tool for patient stratification, we mention some limitations.

Regarding preanalytical phase, our model was fitted for HES-stained tumor slides, and has not been yet validated on HE-stained histopathological slides, despite it being the staining of choice in pathology laboratories outside of France. Adaptations of the model for an optimal application on HE-stained slides and other routine stainings are currently under development. As for the scanning of the slides, only two different scanners were used for the digitization.

The population size of the validation with a majority of patients coming from the same center is also a limitation. On those issues, assorted data from novel external centers is being collected and included in future validation cohorts, which will allow us to confirm the medical utility of RlapsRisk, as recommended in (46).

A prospective observational study comparing RlapsRisk to molecular prognostic signatures is currently ongoing, and will allow us to apprehend the practical benefits and the real impact of the implementation of this AI-based tool into the daily workflow. Future extensions of our research include the development of novel algorithms, adapted to a broader variety of inputs. To deepen interpretability issues, an exhaustive analysis of tiles is contemplated with a focus on the spatiality notion that could provide novel insights in tumor biology.

## CONCLUSION

Our DL model (RlapsRisk) based on single HES-stained tumor WSI showed encouraging performance in the prediction of metastasis free survival among patients with ER+ HER2-localized breast cancers who received state of the art treatments. The use of RlapsRisk score combined with existing clinico-pathological factors improved identification of patients at high risk of relapse, constituting a promising tool for helping treatment decisions. Additionally, identification and characterization of the most contributive parts in the WSI present opportunities to better understand this complex disease and the different features that can reflect a poor or good prognosis.

