## Supplementary material for "Deep Learning Allows Assessment of Risk of Metastatic Relapse from Invasive Breast Cancer Histological Slides"

#### Flow Charts

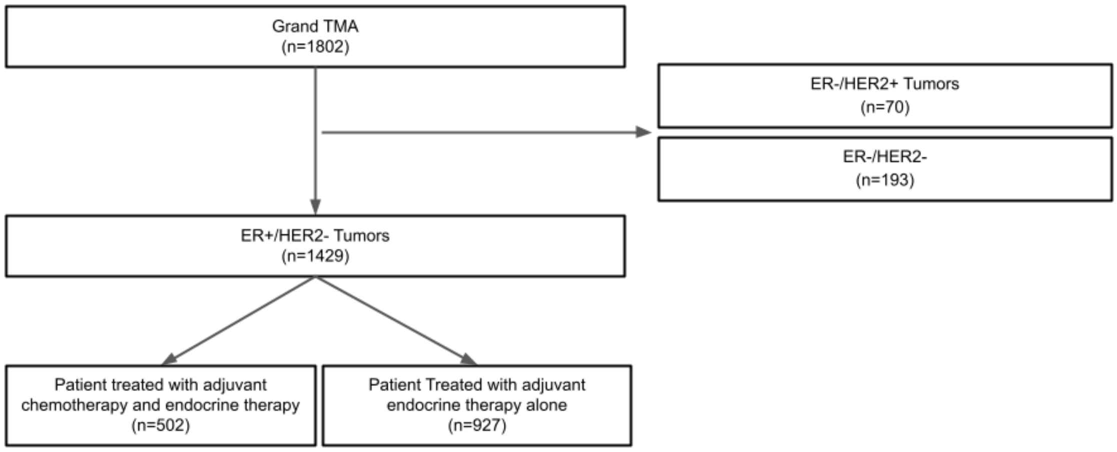


*Supplementary Figure 1: Flow Diagram GrandTMA Cohort*

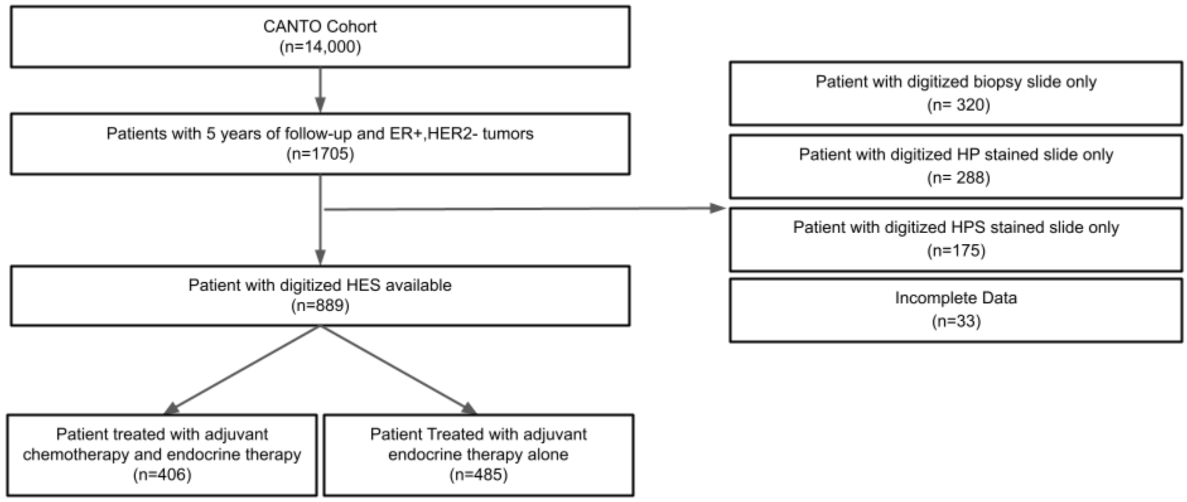


*Supplementary Figure 2: Flow Diagram CANTO Cohort*

| | | TMA | | CANTO | |
|---|---|---|---|---|---|
| **Clinical Variable** | Group | *N* | % | *N* | % |
| **MFS events** | None | **1379** | **92,61%** | **843** | **94,83%** |
| | before 5 years | **57** | **3,83%** | **34** | **3,82%** |
| | after 5 years | **53** | **3,56%** | **12** | **1,35%** |
| **Follow-up** | Mean (sd) | **77 months(41)** | **NA** | **69 months(24)** | **NA** |
| **Age** | Mean (sd) | **61.2 (12.4)** | **NA** | **57.7 (12.2)** | **NA** |
| **Menopausal Status** | Post-menopausal | **1070** | **74.88%** | **667** | **74.86%** |
| | Pre-menopausal | **359** | **25.12%** | **224** | **25.14%** |
| **pT** | 1 | **979** | **68.51%** | **592** | **66,44%** |
| | 2 | **398** | **27.85%** | **265** | **29,74%** |
| | 3 | **50** | **3.40%** | **33** | **3,70%** |
| | 4 | **3** | **0.21%** | **0** | **0,00%** |
| | NA | **2** | **0.14%** | **1** | **0,11%** |
| **pN** | 0 | **950** | **66.48%** | **593** | **66,55%** |
| | 1 | **359** | **25.12%** | **227** | **25,48%** |
| | 2 | **72** | **5.04%** | **46** | **5,16%** |
| | 3 | **48** | **3.36%** | **24** | **2,69%** |
| **Lymph node Invaded** | N0 | **950** | **66.48%** | **753** | **84,51%** |
| | N+ | **479** | **33.52%** | **138** | **15,49%** |
| **Tumor Grade** | G1 | **428** | **29.95%** | **185** | **20,76%** |
| | G2 | **736** | **51.50%** | **537** | **60,27%** |
| | G3 | **263** | **18.40%** | **169** | **18,97%** |
| | NA | **2** | **0.14%** | **0** | **0,00%** |
| **Ki67** | <20% | **1150** | **80,48%** | **756** | **84,85%** |
| | >20% | **279** | **19,52%** | **135** | **15,15%** |
| **Radio** | YES | **1217** | **85.16%** | **475** | **53,31%** |
| | NO | **212** | **14.84%** | **516** | **57,91%** |
| **Hormonoth** | YES | **1387** | **97.06%** | **856** | **96,07%** |

| | | | | | |
|---|---|---|---|---|---|
| | NO | **42** | **2.94%** | **35** | **3,93%** |
| **Chemotherapy** | YES | **505** | **35.34%** | **485** | **54,43%** |
| | NO | **924** | **64.66%** | **406** | **45,57%** |

*Supplementary Table 1 : Patients characteristics in the discovery and validation datasets*

### Integrating a tumor-related feature in the algorithm

To train the tumor classification algorithm, we had at our disposal the 1759 slides of the GrandTMA database whose tumor regions had been contoured by two expert pathologists. The feature extraction pipeline presented in the Model Description was re-used and the multiple instance learning model was replaced by a binary classifier composed of a linear layer with 2048 neurons that classified each tile as tumor vs. non tumor.

#### KM Analyses

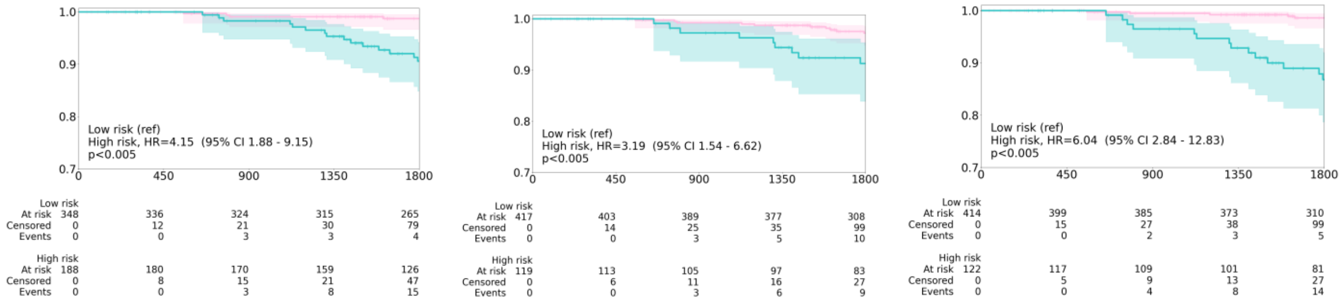


*Supplementary Figure 1: Stratification performed by RlapsRisk Classifier (left) Clinical Score Classifier (middle) Combined Model Classifier (right) on Grade 2 patients from the Canto cohort.*

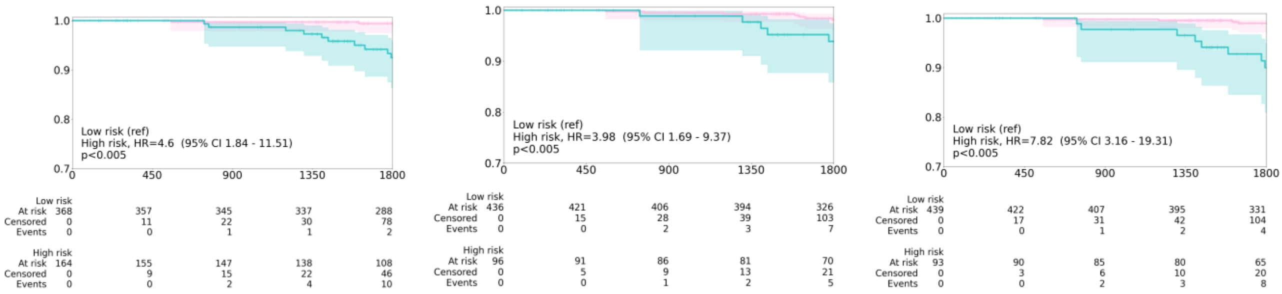


*Supplementary Figure 2: Stratification performed by RlapsRisk Classifier (left) Clinical Score Classifier (middle) Combined Model Classifier (right) on "intermediate risk" patients (defined in Table 1) from the Canto cohort.*

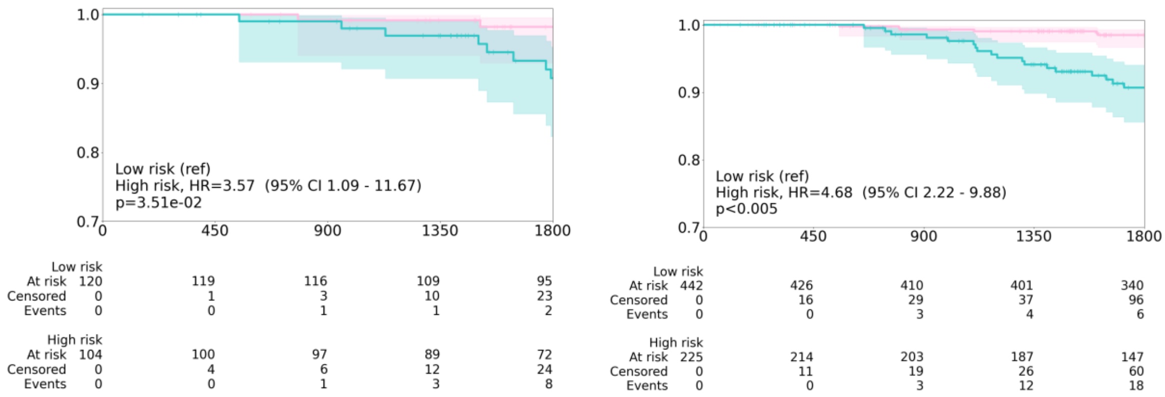


*Supplementary Figure 3A: Stratification performed by RlapsRisk Classifier on pre-menopausal patients of Canto (left), post-menopausal patients (right)*

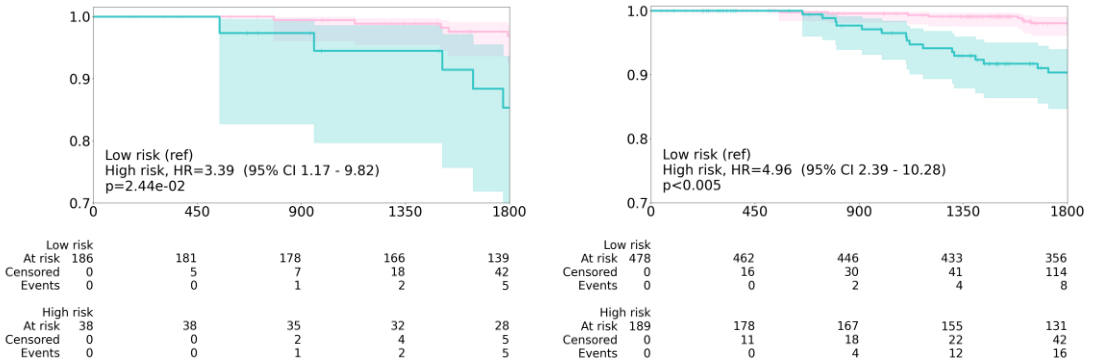


*Supplementary Figure 3B: Stratification performed by the Clinical Score Classifier on pre-menopausal patients of Canto (left), post-menopausal patients (right)*

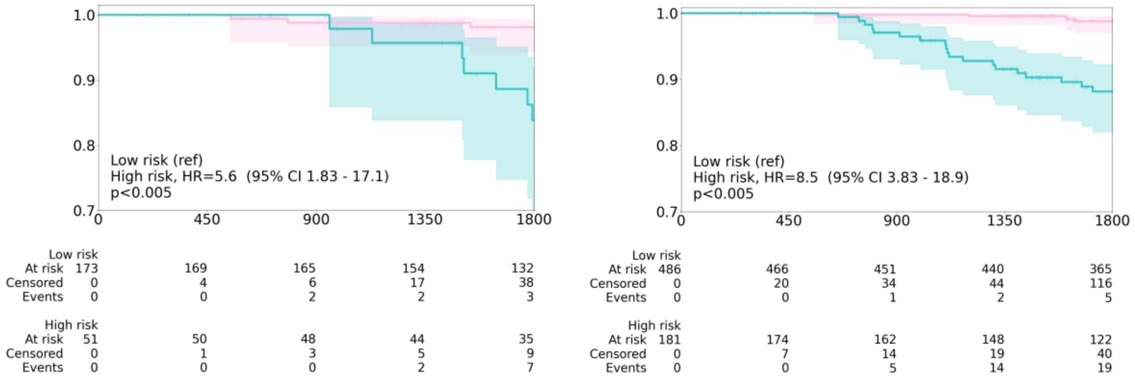


*Supplementary Figure 3C: Stratification performed by the Model Combined Classifier on pre-menopausal patients of Canto (A), post-menopausal patients (B)*

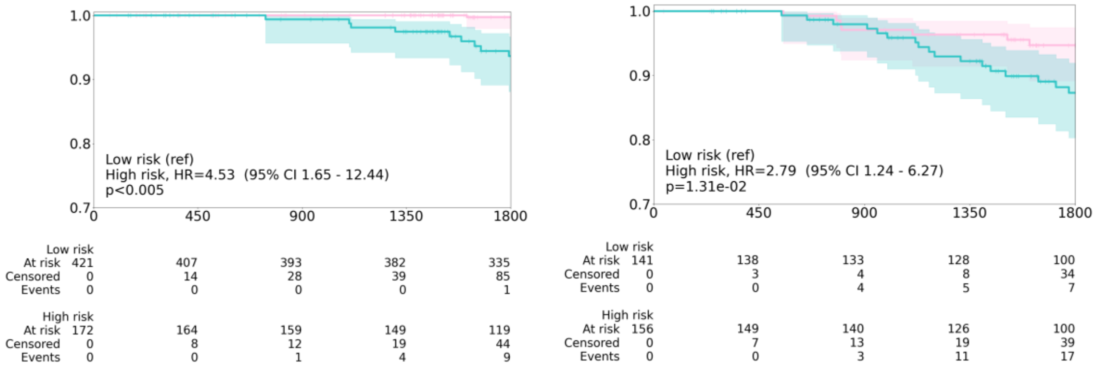


*Supplementary Figure 4A: Stratification performed by RlapsRisk Classifier on patients without lymph node invasion of Canto (left), patients with lymph-node invasion (right).*

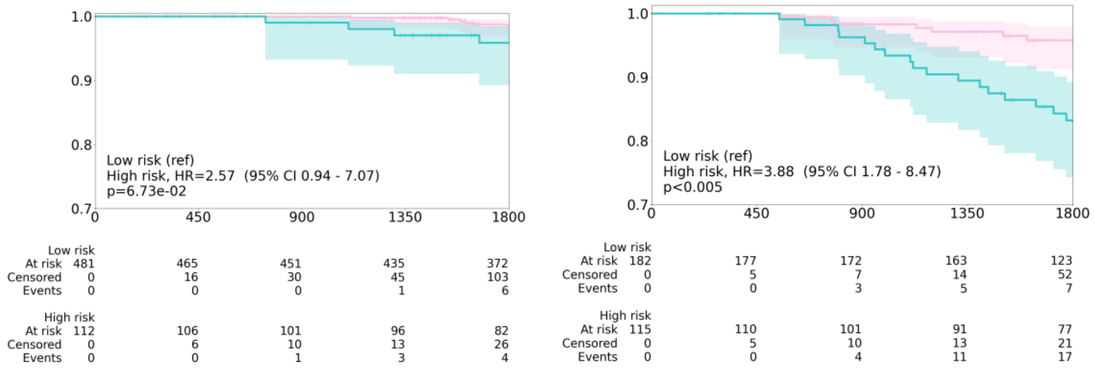


*Supplementary Figure 4B: Stratification performed by the Clinical Score Classifier on patients without lymph node invasion of Canto (left), patients with lymph-node invasion (right).*

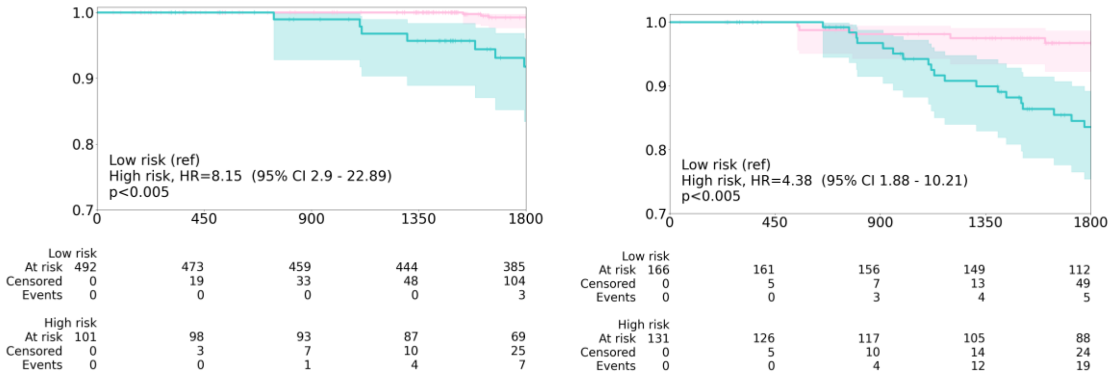


*Supplementary Figure 4C: Stratification performed by the Model Combined Classifier on patients without lymph node invasion of Canto (left), patients with lymph-node invasion (right).*

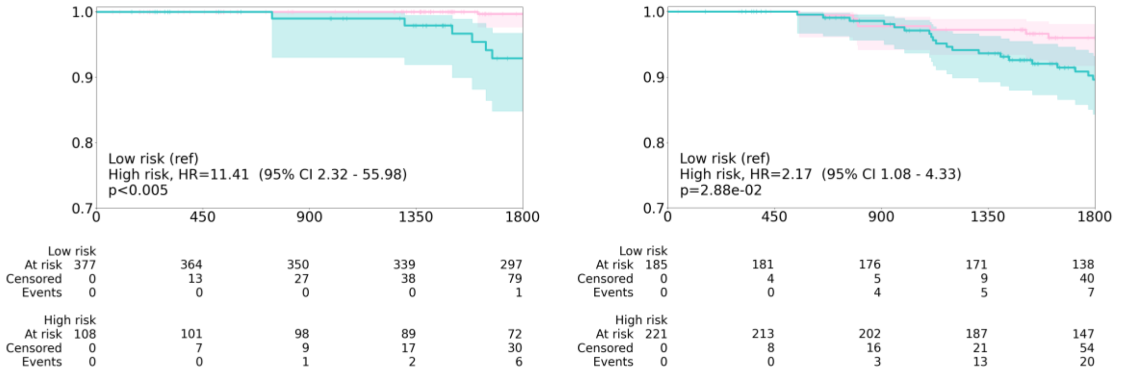


*Supplementary Figure 5A: Stratification performed by RlapsRisk Classifier on patients treated with endocrine therapy alone in Canto (left) , patients treated with chemo-endocrine therapy (right)*

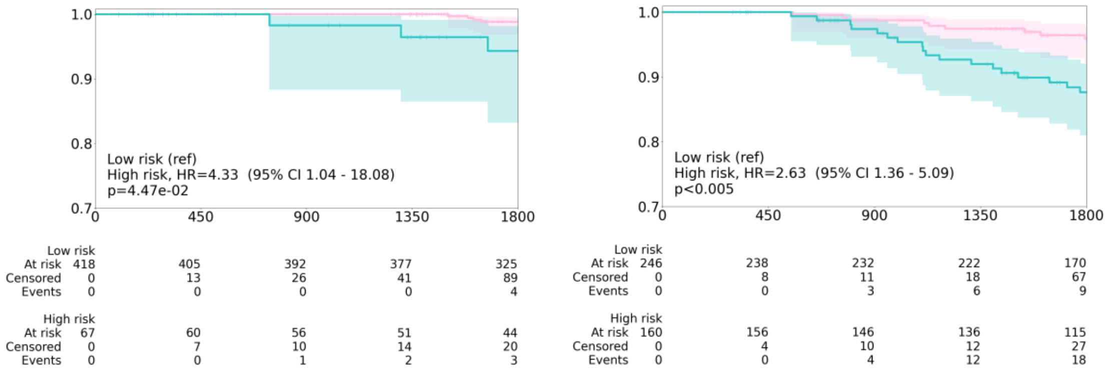


*Supplementary Figure 5B: Stratification performed by the Clinical Score Classifier on patients treated with endocrine therapy alone in Canto (left) , patients treated with chemo-endocrine therapy (right)*

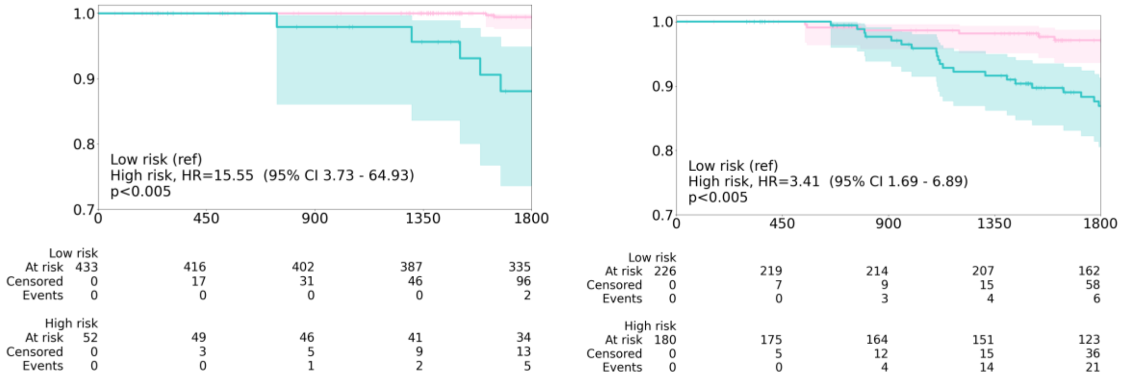


*Supplementary Figure 5C: Stratification performed by the Model Combined Classifier on patients treated with endocrine therapy alone in Canto (left) , patients treated with chemo-endocrine therapy (right)*
